# Multi-sample, multi-platform isoform quantification using empirical Bayes

**DOI:** 10.1101/2025.02.08.637184

**Authors:** Arghamitra Talukder, Shree Thavarekere, Madison Mehlferber, Gloria M Sheynkman, David A. Knowles

## Abstract

Accurate quantification of RNA isoform abundance is crucial for understanding gene regulation, cellular behavior, and disease mechanisms. While short-read (SR) sequencing provides high-throughput and cost-effective transcript quantification, it suffers from read-to-transcript ambiguity. Long-read (LR) sequencing reduces this ambiguity but faces challenges such as high error rates, biases, and lower throughput. Existing methods rely on either SR or LR data and operate on single or merged samples, failing to leverage the variability across multiple samples and the complementary strengths of both technologies. As a result, they struggle to accurately quantify low-abundance and moderate-expressed isoforms and often require complex models for sample-specific bias correction. To address these limitations, we introduce **JOLI**, a hierarchical model that leverages multi-sample learning to enhance transcript quantification by jointly integrating SR and LR sequencing data. By incorporating multi-sample learning, JOLI captures shared transcript structures, corrects for systematic biases, and enhances statistical power, particularly for low- and moderate-abundance isoforms. Our model applies an empirical Bayes framework, learning a shared prior across samples to improve inference consistency. By jointly modeling SR and LR data, it integrates the strengths of both technologies, achieving higher accuracy and reproducibility in transcript quantification. Through benchmarking on simulated and real RNA-seq datasets, we show that JOLI consistently outperforms single-sample EM method by improving ranking consistency, proportional agreement, and estimation accuracy while enhancing reproducibility. Specifically, in simulations, JOLI multi-sample improves Spearman correlation by 9.8% for LR and 7.7% for SR data compared to single-sample method, while for real data, the improvements are 2.56% (LR) and 1.28% (SR), respectively. Multi-sample learning further improves the quantification of isoforms with low to moderate expression levels. Furthermore, JOLI performs competitively with state-of-the-art methods, highlighting its robustness in transcript quantification.

## 1 Introduction

The transcription of DNA into RNA is an intricate process, where individual genes can produce multiple RNA isoforms through mechanisms such as alternative splicing. Accurately quantifying transcript abundance, especially at the isoform level, is fundamental in genomics, shedding light on cellular behavior [29], developmental dynamics [16], and disease mechanisms [10].

Over the past decade, RNA-seq has transformed transcriptomics by enabling high-resolution analysis of gene expression and insights into isoform-level variation. Short-read (SR) sequencing technologies [3], which are cost-effective and extremely high-throughput, enable reproducible quantification. However, its short read length—typically 100-150 base pairs (bp)—is much smaller than the median length of spliced mRNA transcripts (around 3,300bp in humans) [24]. This discrepancy leads to a fundamental issue of read-to-transcript ambiguity, where the origin of many reads cannot be precisely determined. This ambiguity contributes to uncertainty in transcript abundance estimates, which complicates downstream analyses. In contrast, emerging long-read (LR) RNA sequencing technologies, such as those from Pacific Biosciences (PacBio) [6,25] and Oxford Nanopore Technologies (ONT) [20,31], offer reads spanning thousands of base pairs, potentially even covering the entire full-length transcript. This reduces or in the latter case eliminates ambiguity in transcript assignment and facilitates more accurate transcript discovery and quantification. PacBio long reads, especially HiFi reads, are highly regarded for their exceptional accuracy and minimal error rates [11]. Nevertheless, this advantage comes at the expense of reduced throughput and higher costs when compared to ONT sequencing. Conversely, ONT sequencing provides much greater throughput, often making it a more cost-efficient option, but the reads generated typically exhibit lower accuracy and increased error rates ranging from 5 to 20% [12,13]. Indeed, the recent LR-GASP benchmarking effort showed that for isoform quantification (as opposed to discovery), SR analysis currently outperforms LR [21]. This motivates the question: can we get the best of both worlds, combining the throughput and accuracy of SR data with the reduced ambiguity of LR?

Many methods have been developed for isoform quantification for SR platforms— -BitSeqVB [9], RSEM [14], Kallisto [5], Salmon [22]—and LR platforms—LIQA [12], NanoCount [8], Oarfish [13], LR-kallisto [19]. These methods are restricted to a single platform and work with either individual or combined samples, preventing them from capturing variability across multiple samples. Consequently, they are unable to integrate both the distinct advantages of SR and LR technologies for transcript quantification. To address this, we introduce **JOLI**, a hierarchical model for *JOint Long and short Isoform quantification*. JOLI builds upon the standard EM framework by incorporating two key extensions: 1) joint analysis of SR and LR data, and 2) sharing information between multiple similar samples (“multisample” analysis). Our proposed method simultaneously leverages read information from multiple short and long-read samples, exploiting their complementarity to improve accuracy, consistency and reproducibility. It operates on multiple related samples (e.g., from the same tissue type), under the assumption that the true isoform abundances for these samples will be correlated. Applying an empirical Bayes framework, we learn a common prior across the samples that allows the model to capture shared variability and facilitates robust and consistent inference. To estimate the Maximum A Posteriori (MAP) parameters, we employ the EM algorithm within each sample followed by gradient descent for the shared prior parameters.

### Multi-sample variability

Our hierarchical model accounts for sample variability, enabling more accurate estimation of isoform abundances while addressing biases that are not generalizable across different sequencing technologies [18]. SR and LR sequencing technologies are subject to different biases and artifacts, such as PCR amplification and reverse transcription bias [23,1,26]. LR sequencing faces challenges such as 3’ bias, caused by technical limitations like pore blockages, incomplete cDNA synthesis, or RNA degradation. As a consequence, LR can exhibits length biases, with short transcripts being disproportionately represented [2]. In comparison, SR sequencing offers more uniform coverage due to the relatively random distribution of reads across fragmented RNA, making it well-suited to counteract these biases when the two technologies are used together [26]. Methods such as LIQA [12] and Oarfish [13] rely on complicated models for bias and error correction. By leveraging both SR and LR, our model is able to provide a simpler and more robust solution.

### Enhancing Noise-to-Signal Ratio for Low and moderate Isoform Abundance

By pooling information across samples, our method leverages shared transcript structures, increasing statistical power to detect low and moderate abundance isoforms and distinguish genuine signals from noise, even in the presence of technical biases [28,17]. A common strategy for leveraging multiple samples perform isoform assembly and identification using a merged set of all reads, then quantification of these isoforms independently in each sample [27]. This approach however does not share valuable quantitative information between samples. In contrast, our hierarchical model addresses achieves this by treating observations as noisy realizations of a shared prior to balance variability and consistency across related samples.

## 2 Methods

We introduce the key notations used in this work in Table 1 and present the generative model underlying our framework in Fig. 1. This section details the generative model, JOLI model fitting, and dataset.

**Table 1:** Summary of notations.

| Symbol | Description |
| --- | --- |
| $R$ | Number of reads |
| $I$ | Number of isoforms/transcripts |
| $F$ | Number of samples |
| $d_r$ | The $r$-th read |
| $t_i$ | The $i$-th transcript |
| $\alpha$ | Dirichlet parameter of transcript abundance |
| $\theta_{fi}$ | Abundance of transcript $t_i$ in the sample $f$ |
| $a_{ri}$ | True binary indicator if the $d_r$ is from $t_i$. |
| $y_{ri}$ | Assignment matrix $y_{ri} = 1$ if the $d_r$ is consistent with $t_i$, or $y_{ri} = 0$ otherwise |
| $z_{ri}$ | Probability matrix, that the $d_r$ is generated by $t_i$ |
| $n_i$ | Expected number of reads generated by $t_i$. |

**Fig. 1:**
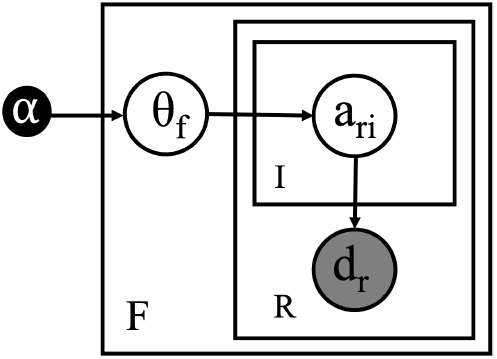
Generative model for multi-sample reads.

**Fig. 2:**
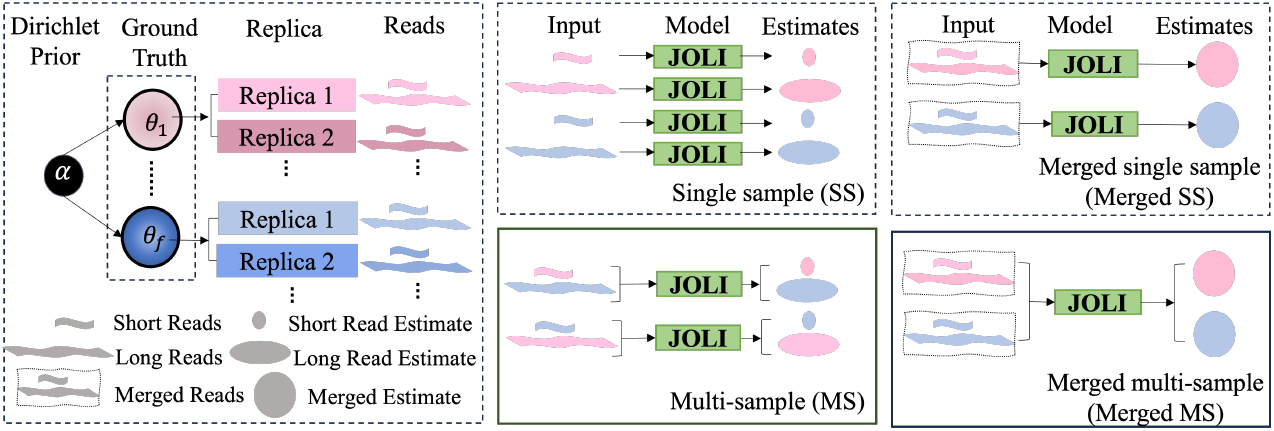
Overview of the experimental design. The leftmost section illustrates the data generation process. Dirichlet prior (*α*) generates multiple ground truth proportions (*θ*_1_ … *θ*_*f*_), each of which produces multiple replicates. These replicates, generate both short and long sequencing reads. The right side of the figure presents the four experimental setups for evaluating the model: *single sample (SS), multi-sample (MS), merged single sample (merged SS)*, and *merged multi-sample (merged MS)*.

### 2.1 JOLI Generative Model

Our generative model describes RNA-seq reads originating from multiple samples and builds upon the single-sample case by incorporating sample-specific parameters (Fig. 1). Each sample *f* has its own transcript proportions *θ*_*f*_, reflecting differences across biological conditions or individuals. Each *θ*_*f*_ is drawn from a Dirichlet distribution, i.e. *θ*_*f*_ |*α ∼* Dirichlet(*α*), where *α* is a hyperparameter vector. The Dirichlet serves as a shared prior over the transcript proportions, encoding shared information across all samples while allowing sample-specific variations. Defining *α*_○_ = Σ _*i*_ *α*_*i*_ the prior mean is *α/α*_○_ and larger values of *α*_○_ lead to more uniform transcript proportions across samples, while smaller values allow for greater variability. For each read *d*_*fr*_ in sample *f*, the generative process is:

1. A transcript *t*_*fr*_ is sampled from a categorical distribution parameterized by *θ*_*f*_, i.e., *t*_*fr*_ |*θ*_*f*_*∼* Categorical(*θ*_*f*_). This step models that the prior probability that a read comes from a specific isoform is proportion to its relative abundance. For convenience, we define the binary variable *a*_*fri*_, where *a*_*fri*_ = 1 if read *d*_*fr*_ originates from transcript *i* (i.e., if *t*_*fr*_ = 1) and *a*_*fri*_ = 0 otherwise. The contribution of this step to the joint probability is,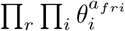.
2. The observed read *d*_*fr*_ is generated conditionally on the selected transcript *t*_*fr*_ according to,

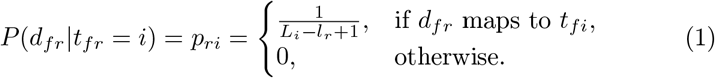

where *l*_*r*_ is the read length and *L*_*i*_ is the transcript length.

The joint probability of the model is,

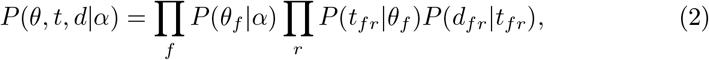

where,

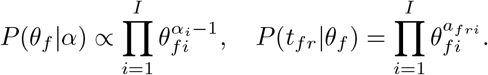

### 2.2 JOLI Model Fitting

We use an expectation-maximization (EM) algorithm to estimate transcript proportions *θ*_*f*_ for each sample *f*, given observed RNA-seq reads *r*. Conditional on the prior parameter *α*, the computations are independent for each sample *f*, so for notational brevity we drop the indexing by *f*. The Dirichlet prior, which governs the distribution of *θ*, is then optimized by leveraging isoform abundance *θ*_*f*_ estimates from all samples. This step serves as the key mechanism for integrating multi-sample information.

#### E: Soft-assigning isoforms to reads

In the E-step we assign reads to isoforms given the current estimate of isoform proportions *θ*. For each read *r* this involves finding the distributions,

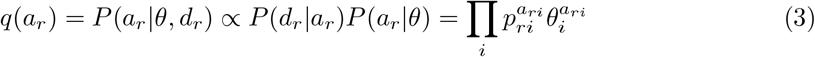

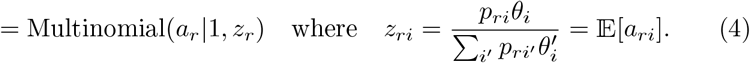

*z*_*ri*_ is our current estimate of the posterior probability that read *r* came from isoform *i*.

#### M: Estimating isoform abundance

For the M-step we need the expected log-likelihood which from Eq. (2) is,

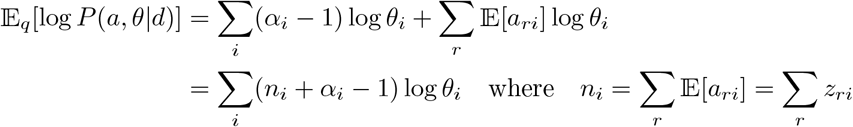

This is the log pdf of a Dirichlet distribution on *θ* with parameter vector *α*^*′*^ = (*α*_1_ +*n*_1_, …, *α*_*I*_ +*n*_*I*_). Standard EM would set *θ* to the mode of this distribution, but we find using the posterior mean 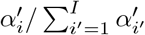 to be more numerically stable.

#### Empirical Bayes prior update

To optimize the Dirichlet hyperparameter vector *α*, we maximize the likelihood of *θ*_*f*_ across all samples, i.e.,

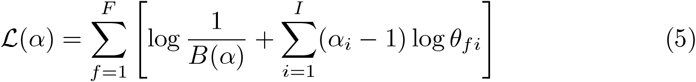

The normalizing constant is *B*(*α*) =Π _*i*_ *Γ* (*α*_*i*_)*/ Γ* (Σ _*i*_ *α*_*i*_). The likelihood in Eq. (5) cannot be optimized analytically so we use gradient ascent to optimze *α*.

In the next iteration, the updated 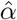 is used in the posterior expectation calculation of *θ*, and this process is repeated iteratively until convergence defined as when the difference between successive estimates, |*θ*^(*s*)^ *−θ*^(*s−*1)^ |, falls below 0.001.

### 2.3 Dataset

#### PacBio and Illumina data

We utilized PacBio HiFi long reads [11] alongside compatible Illumina short reads for our analysis, as illustrated in Fig. 7. The dataset was collected across various stages (days 0–5) of the differentiation protocol for WTC11-derived primordial endothelial cells, with 2 biological replicates for each day. For multi-sample models, we specifically focused on data from day 3 and day 5. Long reads were aligned to the human transcriptome using `Minimap2 [v2.17]` [15], while short reads were processed with `STAR [v2.5.2a]`[7]. To resemble a more typical LR coverage, the LR alignment files were downsampled to 10% of the original reads. Post-downsampling, the dataset consisted of approximately 100 million short reads and 10 million long reads for each sample, originating from around 150,000 transcripts.

#### Simulated data

Using the LRGASP-simulation pipeline [21], two samples, each containing PacBio and Illumina reads, were simulated. PacBio reads were generated using `IsoSeqSim [v0.1]` [30], while Illumina reads were simulated with `RSEM [v1.3.0]` [14]. The entire human genome served as the reference for this simulation. Transcript abundance estimates from real PacBio reads (as described in the previous section) were used to create the expression profile. Each simulated sample consisted of 10 million short reads and 1 million long reads. The ground truth isoform abundance file, obtained from the human genome, transcriptome, and annotation files using the LRGASP pipeline, contained approximately 149,500 isoforms per sample.

## 3 Results

To understand how multi-sample learning provides advantages over single-sample processing, we evaluate our model in four different setups as shown in (Fig. 7): *SS (Single Sample), MS (Multi-Sample), Merged SS (Merged Single Sample)*, and *Merged MS (Merged Multi-Sample)*. Each setup varies in how read data is provided to the model and how abundance estimates are computed.

In SS baseline, each sample is treated completely independently without any information sharing prior optimization. In MS, one LR and one SR dataset, are treated as unique samples (with their own *θ*_*f*_), while still being part of a unified analysis with a shared Dirichlet prior. This allows us to assess the benefits of multi-sample variability learning over single-sample estimation. In merged SS, SR and LR from the same biological replicate are combined into a single “sample” before being processed through EM without any prior optimization, giving a single abundance estimate per ground truth proportion (*θ*), allowing the model to integrate complementary information from different read types. Finally, merged MS integrates both sample merging and multi-sample learning. This configuration evaluates whether aggregating information across multiple samples improves the robustness of abundance estimation, even after merging short and long reads within each sample. For simulation data, all four setups were evaluated, whereas for real data, only SS and MS were conducted.

To comprehensively evaluate the accuracy and reliability of isoform quantification, we employ four metrics: Spearman correlation (SC), Pearson correlation (PC), Median Relative Difference (MRD), and Irreproducibility Measure (IM); all definitions can be found in Appendix 6. Each of these metrics captures a distinct aspect of model performance, ensuring a balanced assessment. SC measures rank-based association, making it robust to non-linear scaling and outliers, while PC captures linear relationships, ensuring proportionality with true expression levels or replicates. Higher values of both indicate better performance. For simulated data where true abundances are known, MRD measures the relative deviation of estimated values from the ground truth, capturing error magnitude. Finally, IM measures transcript abundance irreproducibility across replicates in real data. It is computed based on the coefficient of variation (CV) of transcript abundances. Lower MRD indicates better approximation of true expression levels, while lower IM signifies higher reproducibility. For simulation data, evaluation metrics were computed with respect to the ground truth, while for real data, they were assessed based on replicates.

### Simulations results

Fig. 3 compares estimated transcript abundances log_*e*_(TPM + 1) against ground truth across different evaluation setups. Comparing SS and MS setups, we observe that MS learning consistently improves performance. For long reads, SC and PC improve by approximately 9.8% and 9.9% (Fig. 3, left). For short reads, SC increases by 7.7% and PC by 0.4%(Fig. 3, center). The merged setup, combining long and short reads per sample, shows SC improving by 2.9% and PC by 0.2% (Fig. 3, right).

**Fig. 3:**
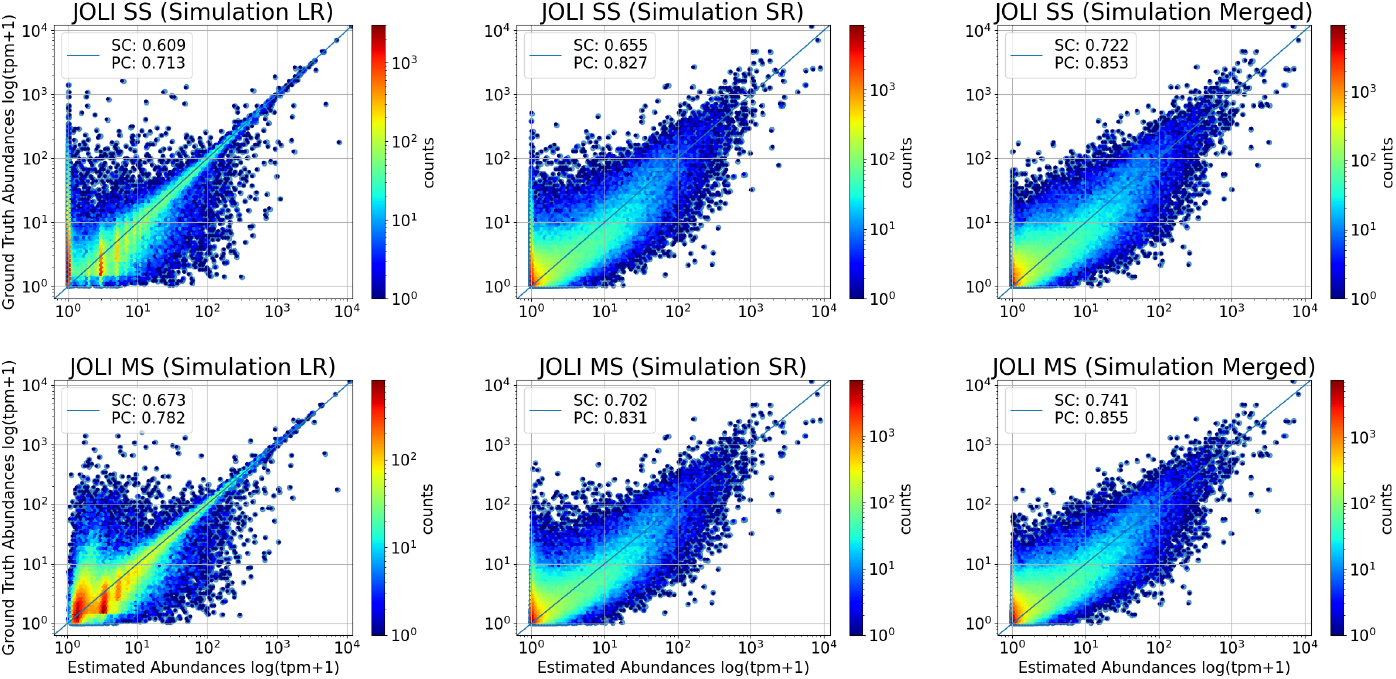
Scatter plots of estimated vs. ground truth transcript abundances. (log_*e*_(TPM+ 1)) for the simulation dataset under different experimental setups. Multi-sample learning improves abundance estimation, with higher SC and PC, demonstrating the benefits of leveraging cross-sample information.

Fig. 4 highlights the contribution of MS learning in enhancing the noise-to-signal ratio for low- and moderate-expressed isoforms by assessing SC, PC, and MRD across three transcript expression categories—low, moderate, and high. The classification is based on ground truth abundance, with varying mean and standard deviation detailed in Table 2. Correlation is particularly enhanced for low and moderate expressed transcripts. For LR (Fig. 4, left), JOLI MS outperforms JOLI SS at moderate expression levels in SC and PC but struggles at low levels with a higher MRD. For SR (Fig. 4, center), JOLI MS improves MRD across all expression levels and outperforms JOLI SS in SC and PC for low and moderate expression. In merged experiments (Fig. 4, right), JOLI MS follows the same trend as SR, exhibiting higher SC and PC at both low and moderate expression levels, while consistently achieving better MRD across all expression ranges..

**Table 2:** Number of isoforms, average abundance (log_*e*_(TPM+ 1)), and standard deviation across different abundance classes.

| Expression Level | # of Isoforms | Avg. Abundance | Std Dev |
| --- | --- | --- | --- |
| Low | 50,531 | 0.087 | 0.055 |
| Moderate | 50,731 | 0.479 | 0.185 |
| High | 49,831 | 1.958 | 1.076 |

**Fig. 4:**
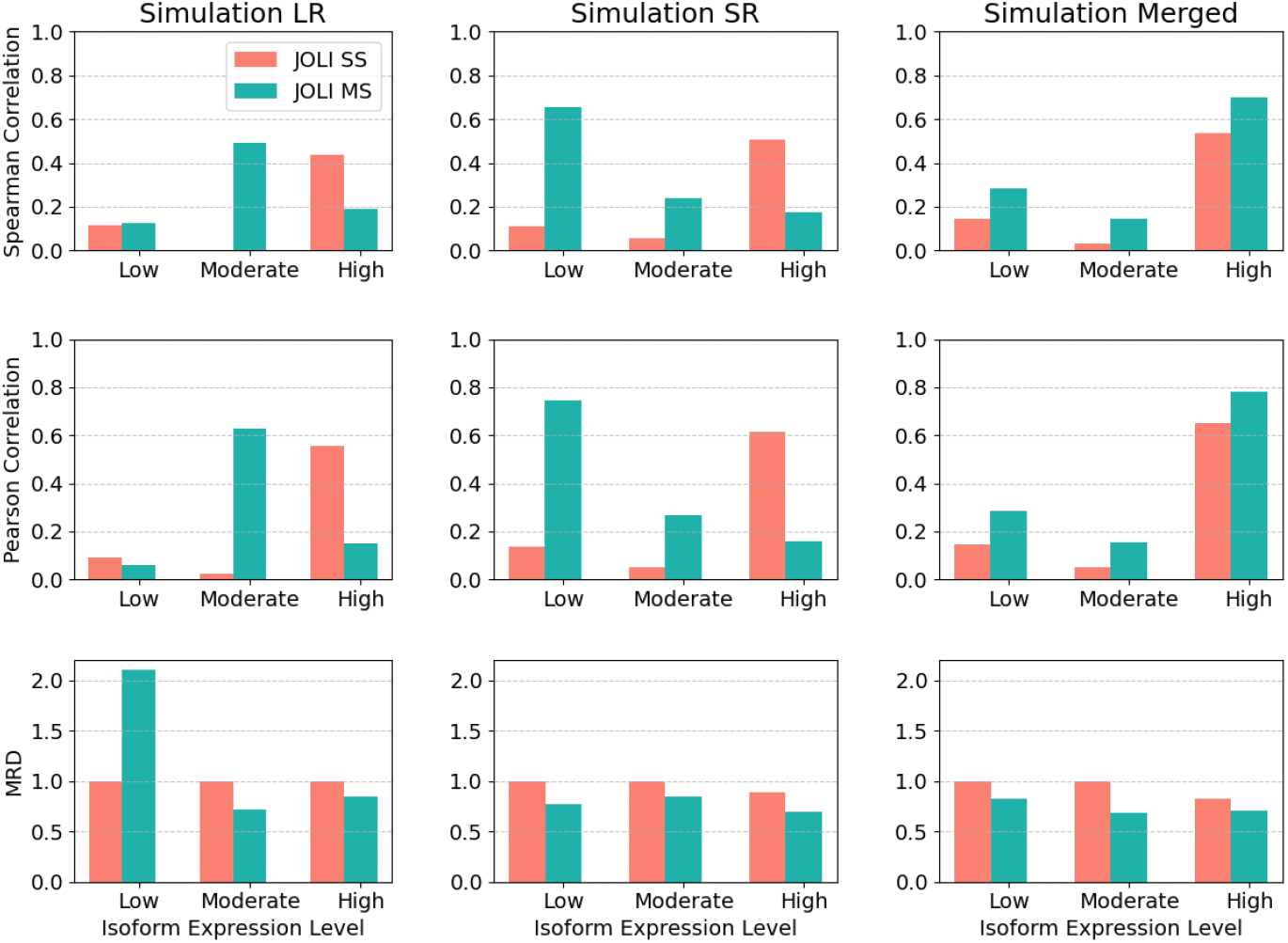
**SC, PC, and MRD comparisons for low, moderate, and highly expressed isoforms** in simulated data. For LR (left), JOLI MS outperforms JOLI SS in SC and PC at moderate expression but has higher MRD at low levels; for SR (center), JOLI MS improves MRD across all levels and outperforms JOLI SS in SC and PC at low and moderate expression; in merged experiments (right), JOLI MS shows higher SC and PC at low and moderate expression with consistently better MRD at low and moderate levels.

Fig. 5a showcases a performance comparison of JOLI SS and MS against state-of-the-art (SOTA) methods, including `lr-Kallisto [v0.51.1]` and `Oarfish [v0.6.5]` for LR, as well as `Kallisto [v0.51.1]` for SR. The commands used to run the SOTA methods can be found in Appendix 6. For LR in Fig. 5a (left), the lower SC (0.67) and PC (0.78) for JOLI MS compared to lr-kallisto (0.85, 0.93) and Oarfish (0.81, 0.89) indicate that while JOLI MS captures overall isoform abundance, it struggles with precise ranking and consistency in absolute expression values, likely due to increased variance in MS learning. The most notable advantage of JOLI MS is in MRD, where it achieves 0.61, significantly outperforming JOLI SS and Oarfish (1.0) and approaching lr-kallisto (0.33). This highlights that despite lower SC and PC, JOLI MS reduces systematic biases found in single-sample methods and improves reliability in abundance estimates by mitigating quantification errors. On the other hand, JOLI MS demonstrates a much more competitive performance in SR results Fig. 5a (right). It outperforms kallisto in PC (0.83 vs. 0.78), a slightly lower SC (0.70 vs. 0.72), and a comparable MRD (0.78 vs. 0.77). These results indicate that JOLI MS effectively preserves absolute expression values while maintaining ranking accuracy and quantification error at levels similar to kallisto.

**Fig. 5:**
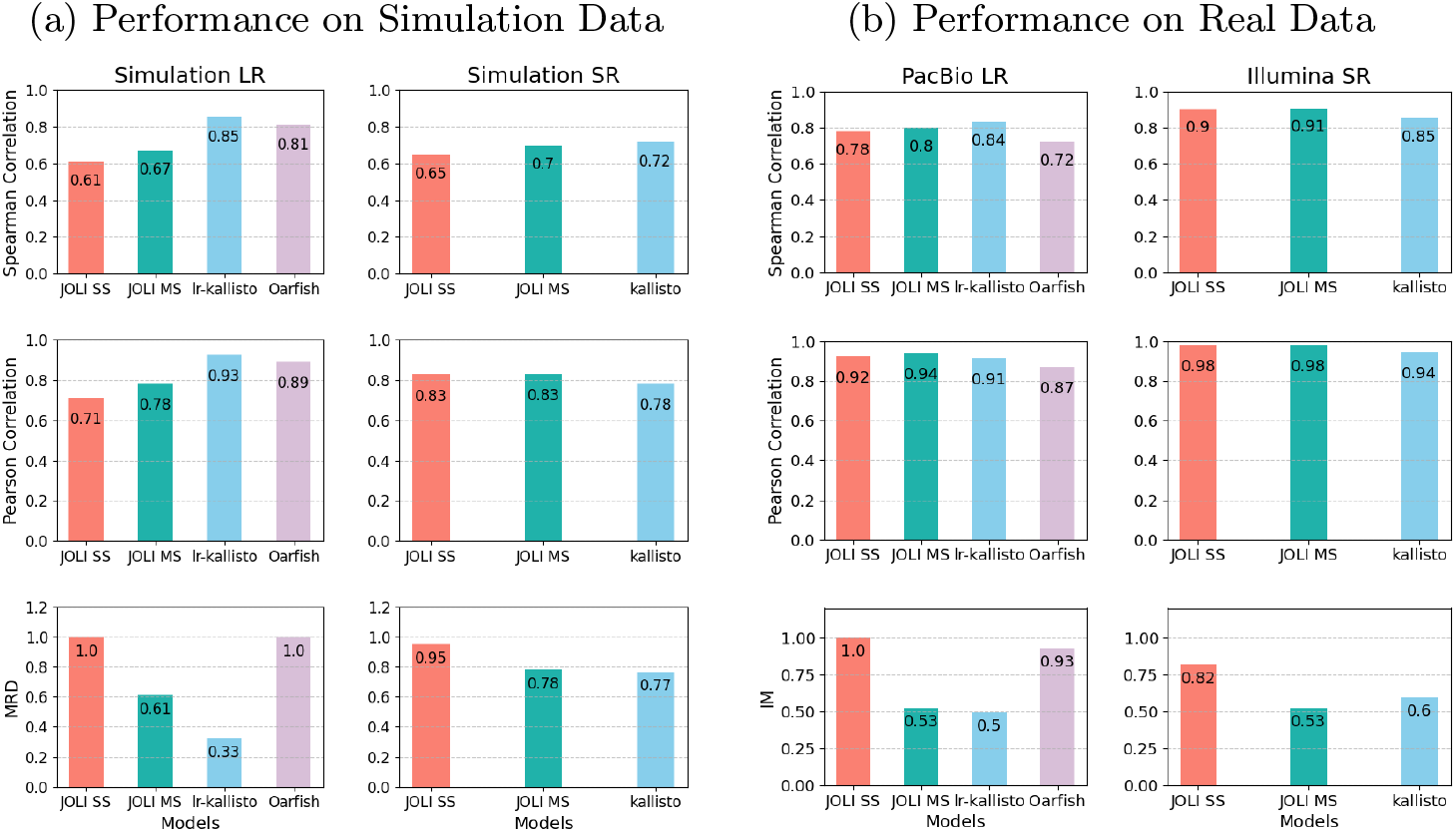
Performance comparison of JOLI models with SOTA for both simulation and real data. (a) Simulation results compare JOLI MS and JOLI SS with lr-kallisto and Oarfish for LR and kallisto for SR across SC (top), PC (middle), and MRD (bottom), showing that JOLI MS reduces MRD while trailing lr-kallisto and Oarfish in SC and PC. (b) Real data results for PacBio LR and Illumina SR indicate that JOLI MS improves over JOLI SS in all metrics, reducing IM while maintaining high correlation, performing similarly to lr-kallisto and outperforming Oarfish. For SR, JOLI MS competes with kallisto, achieving higher SC, comparable PC, and lower IM, emphasizing its reproducibility and ranking consistency.

**Fig. 6:**
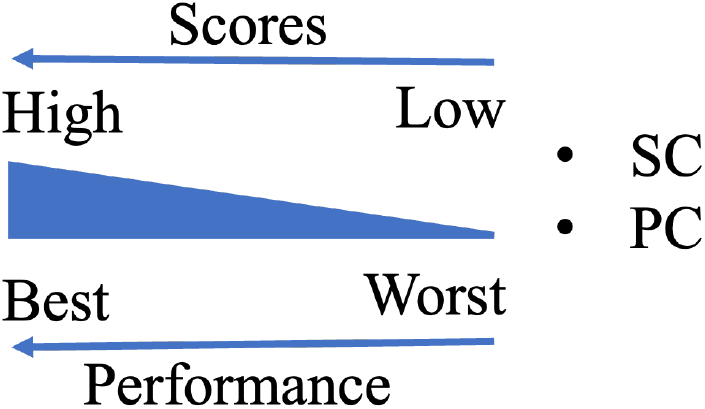
For SC and PC **higher** values mean **better** performance

**Fig. 7:**
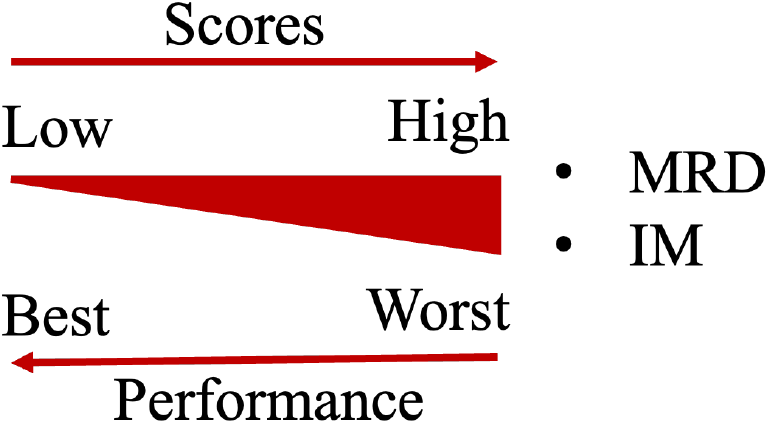
For MRD and IM **lower** values mean **better** performance

### PacBio and Illumina results

Fig. 5b presents a performance comparison of JOLI for PacBio LR and Illumina SR. Fig. 5b (left) shows that both for LR and SR, JOLI MS improves over JOLI SS across all metrics. The SC of JOLI MS (0.80) is slightly lower than lr-kallisto (0.84) but higher than Oarfish (0.72), indicating that MS learning enhances rank preservation. PC remains consistently high across methods, with JOLI MS (0.94) performing on par with JOLI SS and lr-kallisto (0.92). The most significant improvement is seen in the IM, where JOLI MS (0.53) substantially reduces irreproducibility compared to JOLI SS (1.0) and even outperforms Oarfish (0.93), though lr-kallisto (0.50) achieves the lowest IM. This suggests that JOLI MS improves reproducibility while maintaining high correlation with the estimated transcript abundances. For SR, Fig. 5b (right) shows JOLI MS performing competitively with kallisto, achieving higher SC (0.91 vs. 0.85), comparable PC (0.98 vs. 0.96), and lower IM (0.53 vs. 0.60), while outperforming JOLI SS across all metrics. Overall, these results demonstrate that JOLI MS provides notable improvements in reproducibility and ranking consistency, making it a strong contender in real-data settings despite minor differences.

## 4 Discussion

We have introduced JOLI, a novel hierarchical model for isoform quantification that jointly analyzes SR and LR sequencing data from multiple samples. Our approach extends the standard EM framework by incorporating information sharing across multiple related samples through an empirical Bayes framework which learns a common prior to capture shared variability. By integrating both sequencing technologies, JOLI aims to combine the accuracy and throughput of SR data with the reduced ambiguity of LR data. During the development of JOLI, another joint model, MPAQT [4], was proposed to integrate LR and SR. However, it does not model multiple samples, relies on precomputed LR abundance estimates, requires memory-intensive simulated data, and assumes unambiguous LR alignment.

Through benchmarking on both simulated and real datasets, we have demonstrated that JOLI MS outperforms its SS counterpart, particularly in reducing estimation errors and improving reproducibility. In simulations, JOLI MS consistently outperforms JOLI SS across different sequencing setups. Further analysis indicates that MS learning has the most significant impact on low- and moderate-abundance transcripts, which are typically harder to quantify. Comparison with SOTA methods shows that JOLI MS achieves competitive results, particularly in error reduction, highlighting its ability to generate more reliable abundance estimates. On real datasets, JOLI MS continues to show strong performance improvements over JOLI SS, demonstrating the practical advantages of MS learning in handling biological variability and noise, while performing at the same level as other state-of-the-art methods.

Our findings demonstrate that MS learning is particularly effective in improving reproducibility and mitigating systematic biases, making it an essential component of accurate isoform quantification. This motivates us to integrate joint SR and LR, and MS, learning into existing state-of-the-art methods to fully leverage its benefits.

## Code Availability

The source code is available at https://github.com/ArghamitraT/JOLI. This code is based on https://github.com/a-slide/NanoCount.

## 5 Acknowledgements

We thank PacBio for providing sequencing data. This work was supported by National Science Foundation (NSF) CAREER DBI2146398 to D.A.K, an NSF CSGrad4US award and NIH Training Grant 5T15LM007079-34 to A.T. and National Institutes of Health (NIH) grant R35GM142647 to G.M.S. Any opinions, findings, and conclusions or recommendations expressed in this material are those of the authors and do not necessarily reflect the views of the NSF or the NIH.

## 6 Appendix

### 6.1 Performance Evaluation Metric

#### Spearman Correlation (SC)

We denote *θ*_*i*_ as the estimated abundance and 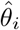 as the ground truth abundance of transcript *i* (*i* = 1, 2, …, *I*). The Spearman correlation between log(*θ* + 1) and 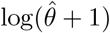 is given by:

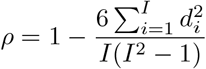

where

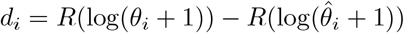

represents the difference between the ranks of the log-transformed estimated abundance *θ* and ground truth 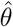.

#### Pearson Correlation (PC)

We denote *θ*_*i*_ as the estimated abundance and 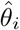 as the ground truth abundance of transcript *i* (*i* = 1, 2, …, *I*). The Pearson correlation coefficient between log(*θ* + 1) and 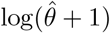 is given by:

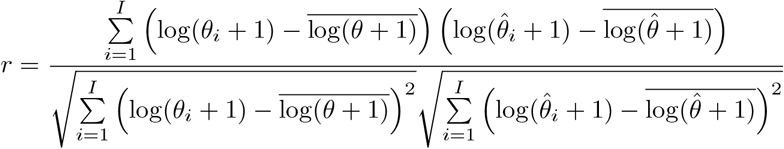

where

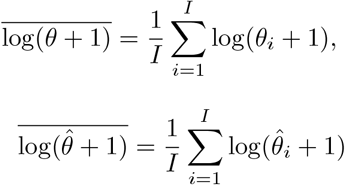

represent the mean of the log-transformed estimated and true abundance values, respectively.

#### Median Relative Difference (MRD)

The MRD measures the median percentage of estimated values that deviate from the true ones:

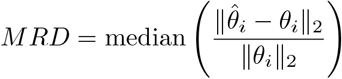

#### Irreproducibility Measure (IM)

Denote *θ*_*ijk*_ as the abundance estimation of transcript (*i* = 1, 2, …, *l*) in a sample, where *j*(*j* = 1, 2, …, *J*) represents different groups (i.e., conditions or tissues) and *k*(*k* = 1, 2, …, *K*) represents different replicates within the group.

Coefficient of variation **CV**_**ij**_ of log *θ*_*ijk*_ (*k* = 1, 2, …, *K*)

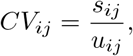

where *s*_*ij*_ and *u*_*ij*_ are the sample standard deviation and mean of abundance estimates.

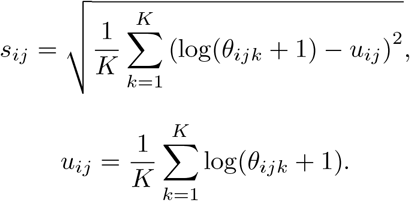

#### Irreproducibility measure based on CV_ij_

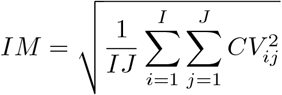

### 6.2 Commands to run SOTA

#### lr-Kallisto

~~~
kallisto bus -x bulk --threshold 0.8 -t 32 --long --unmapped
-i index_file reads_file -o output_dir
bustools sort -t 30 output_dir/output.bus -o sorted.bus
bustools count sorted.bus -t transcripts.txt -e matrix.ec -o count
--cm -m -g t2g.txt
kallisto quant-tcc -t 32 --long -P PacBio count.mtx
-i index_file -e count.ec.txt -o output_dir
~~~

#### Kallisto

~~~
kallisto bus -x bulk --threshold 0.8 -t 32 --paired
-i index_file reads_file1 reads_file2 -o output_dir
bustools sort -t 30 output_dir/output.bus -o sorted.bus
bustools count sorted.bus -t transcripts.txt -e matrix.ec -o count
--cm -m -g t2g.txt
kallisto quant-tcc -t 32 count.mtx
-i index_file -e count.ec.txt -o output_dir
~~~

#### Oarfish

Reads were aligned using `Minimap2 [v2.28].`

~~~
minimap2 --eqx -N 100 -ax map-pb transcripts.fasta reads.fasta
| samtools view -@4 -b -o aln.bam
oarfish -j 16 -a aln.bam -o output_dir/ --filter-group no-filters --model-coverage
~~~

